# Characterization of the olfactory system in *Apis dorsata*, an Asian honey bee

**DOI:** 10.1101/420968

**Authors:** Sandhya Mogily, Meenakshi VijayKumar, Sunil Kumar Sethy, Joby Joseph

## Abstract

The European honeybee, *Apis mellifera* is the most common insect model system for studying learning and memory. We establish that the olfactory system of *Apis dorsata*, an Asian species of honeybee as an equivalent model to *Apis mellifera* to study physiology underlying learning and memory. We created an Atlas of the antennal lobe and counted the number of glomeruli in the antennal lobe of *Apis dorsata* to be around 165 which is similar to the number in the other honey bee species *Apis mellifera* and *Apis florea*. *Apis dorsata* was found to have five antenno-cerebral tracts namely mACT, lACT and 3 mlACTS which appear identical to *Apis mellifera*. Intracellular recording showed that the antennal lobe interneurons exhibit temporally patterned odor-cell specific responses. The neuritis of Kenyon cells with cell bodies located in a neighborhood in calyx retain their relative neighborhoods in the peduncle and alpha lobe forming a columnar organization in the mushroom body. Alpha lobe and the calyx of the mushroom body were innervated by extrinsic neurons with cell bodies in the lateral protocerebrum. A set of GABA positive cells in the lateral protocerebrum send their neurites towards alpha-lobe. *Apis dorsata* was amenable to olfactory conditioning and showed good learning and memory retention at 24 hours. They were amenable to massed and spaced conditioning and could distinguish trained odor from an untrained novel odor.

## Introduction

Honey bees were described as magic well for discoveries in biology by Karl Von Frisch. The European honey bee, *Apis mellifera* is well established as a model system to investigate various fundamental scientific questions at the behavioral, neural and molecular levels. The olfactory conditioning in bees is extensively used for research in learning and memory (Menzel and Erber 1978; Menzel et al. 1993; Menzel and Muller 1996; Giurfa 2007) as features and mechanisms of learning and memory in bees are found to be similar to those in mammals and humans.

A large part of these results has come from studies carried out using the olfactory system of *Apis mellifera* as the model system. In *Apis mellifera* the odor molecules are detected by around 60,000 olfactory receptor neurons (ORN) present in sensilla located on the antennae (Esslen and Kaisling 1976; Kropf et al. 2014). ORNs from one side innervate a single corresponding antennal lobe (AL), the primary olfactory center, through the T1-4 tracts of the antennal nerve (AN) (Suzuki 1975; Mobbs 1982; Galizia et al. 1999; Abel et al. 2001; Krischner et al. 2006). In the AL of *Apis mellifera*, ORNs synapse on to around 800 Projection neurons (PN) and around 4000 local neurons (LN) in dense spheroidal structures called glomeruli which are the morpho-functional unit of the AL (Hildebrand and Shepherd 1997; Anton and Homberg 1999; Hansson and Anton 2000). PNs in Apis *mellifera* connect to the higher olfactory centers the mushroom bodies (MB) through 5 antenno-cerebral tracts (ACT) where they synapse with around 1,80,000 Kenyon cells (KC) (Mobbs 1982; Abel et al. 2001; Krischner et al. 2006; Muller et al. 2002; Rossler and Brill 2013; Zwaka et al. 2016). While the dendrites of the KCs innervate the calyces, the axons project frontally and form the mushroom body peduncle. Subsequently, the axons of KCs bifurcate and the branches innervate the α-lobe and the β-lobe. Olfactory input is received by the lip region and inner half of the basal ring of the MB calyces (Mobbs 1982).

ORNs have odor and concentration dependent response profiles. ORNs that express similar receptor types innervate a single glomerulus (Mombaerts 1996; Rossler et al. 1999; Galizia and Menzel 2000; Carlsson et al. 2002). Axons of the ORNs innervate the cortex whereas PNs and LNs innervate the core of the glomeruli (Pareto 1972; Galizzia et al 1999). In the AL, odor information from the ORNs is processed in the glomeruli, where they evoke odor-cell specific temporally patterned responses or the spatio-temporal odor code (Laurent 1997, Galizia and Menzel 2000). These responses are characteristic for each kind of odorant and highly reproducible (Stopfer et al. 2003; Galizzia et al. 1999; Carlsson et al. 2002). To understand this olfactory code it is essential to understand the glomerular arrangement.

Though in insects the glomerular number, shape and arrangement varies from species to species, their size, shape and location have been reported to be similar across individuals of same species and sex (Rospars 1988). This makes it possible to make anatomical maps of glomeruli for a species to study the odor code. In most species that we know of, each glomerulus receives input from all ORNs expressing a single receptor type. So the count of glomeruli can act as a constraint in searching for receptor genes in the DNA sequence database (Karpe et al. 2016).

Among honeybee species, the glomerular atlas is available for *Apis mellifera* (Arnold et al. 1985; Flanagan and Mercer 1989; Galizia et al. 1999). Among the nine species in the genus *Apis*, in India, the species *Apis cerena, Apis florea and Apis dorsata* are present widely. A comparative study on the structural differences in the olfactory system of drones of Apis *florea* and *Apis mellifera* (Brockman and Bruckner 2001) is available. *Apis florea* is found to have a similar number of glomeruli and olfactory receptor genes compared to *Apis mellifera* (Karpe et al. 2016). PER assays for olfactory learning were effectively tested in *Apis florea* (Kaspi and Shafir 2013) and *Apis cerana* (Wang and Tan 2014; Jung et al. 2017).

In our study we characterized the olfactory system of *Apis dorsata* worker in terms of anatomy, physiology and behavior. We compared the glomerular organization of *Apis dorsata* to that of *Apis mellifera* and created a digital atlas of the glomeruli. We studied the GABA positive innervations of the AL and the extrinsic neurons of the mushroom body. The nature of the antenno-cerebral tracts and the arrangement of the Kenyon cells and tracts were characterized. We show that the relative arrangement of the cell bodies of Kenyon cells in the mushroom body is carried through in the arrangement of axonal fibers in the peduncle and the lobes forming parallel compartments. We further recorded intracellularly from the AL of *Apis dorsata* to show that the AL interneurons respond in odor-cell specific temporal pattern to odor stimuli. We found *Apis dorsata* to be suitable for olfactory PER conditioning

## Materials and Methods

Honey Bee workers (*Apis dorsata*) were collected from the hives in the university campus, cooled in a refrigerator at 4°c and mounted in plastic tubes using adhesive tape. For all the experiments requiring recordings and dye fills the head was immobilized with paraffin wax. For anterograde fills from antennal nerve, the scapus of an antenna was cut and a crystal of dye Dextran Biotin, 3000MW, lysine fixable (BDA 3000; Molecular Probes) or Dextran tetramethylrhodamine, 3000MW, anionic lysine fixable (D3308; Molecular Probes) was inserted using a pulled glass capillary. The cut antenna was sealed with Vaseline to prevent desiccation. The animal was kept in a moist chamber overnight to allow transport of the dye. The next day the brain was dissected out and fixed in 4% PFA for 24 hours. To visualize the efferent tracts of the AL and the MB the cuticle was cut open to expose the brain. After removing trachea and glands a glass electrode containing dye (dextran biotin/dextran tetramethylrhodamine) was inserted into the location of injection and left for a few seconds. The brain was later covered with the piece of cuticle that was previously cut and kept for 3-4 hours before dissection and fixation.

The fixed brains were rinsed thrice in PBS (20 mins each). If dextran biotin was used then the brains were washed in 3% Triton X in PBS and incubated in Streptavidin Alexa Fluor 633 conjugate (S21375, Invitrogen) in 3% Triton X in PBS for 4 days. In those cases where anti-GABA antibody has been used, the brains were incubated in anti-GABA antibody (Sigma A2052) for four days followed by incubation in anti-rabbit IgG secondary antibody conjugated with Alexa Fluor 488 (Invitrogen, A-11008) for four days. In the case of antibodies the PBS washes were for 4 days each. After rinsing thrice in PBS for 20 mins they were dehydrated in ascending ethanol series and mounted on concavity glass slides in methyl salicylate for imaging.

Processed brains were scanned with a laser scanning confocal microscope (Leica TCS SP2, Leica Microsystems or Karl Zeiss LSCM NLO 710, Germany). An excitation wavelength of 568nm was used for dextran rhodamine, 488nm was used for Alexa Fluor 488 and 633nm was used for Alexa Fluor 633. Oil immersion objective lens with 0.80mm working distance was used. Images were acquired at a resolution of either 1024x1024 or 512x512 pixels with 10 X or 20 X objectives.

## Image processing

Out of the many preparations (n=49), the ones in which all the glomerular margins were clearly visible were scanned and out of these three were selected for counting the glomeruli and comparing the glomerular architecture with the published *Apis mellifera* atlases (Arnold et al. 1985, Flanagan and Mercer 1989, Galizia et al. 1999, Krischner et al. 2006). The images were processed using ImageJ (Life line Fiji version 5.1). The outlines of the glomeruli were traced using segmentation editor. Each glomerulus was individually marked by drawing with hand in a few sections and rest by interpolation. The tracts were also segmented for visualization in the same way. A 3D reconstruction of the AL was made using ImageJ. To create the digital atlas, alternate sections of the image stack were transferred to PowerPoint and each glomerulus was labeled following Galizia 1999.

## Electrophysiology

Bees were collected and mounted as described above. Paraffin wax was used to make a cup around the head to hold the saline. The cuticle was cut open exposing the brain. Glands and sheath were removed carefully and the brain was perfused with bee saline (37 mM NaCl, 2.7 mM KCl, 8 mM Na2HPO4, 1.4 mM KH2PO4, Ph 7.2) (Krischner et al. 2006). Local field potential (LFP) was recorded from Mushroom body calyx by placing saline filled blunt glass microelectode with an impedance around 5 MΩ, and amplified using Axoclamp 900A (Molecular Devices). A chlorided-silver wire placed in saline served as the ground. Recording from antennal lobe neurons was made using sharp microelectrodes (70-100MΩ) filled with 200 mM KCl. Microelectrodes were pulled using horizontal puller (Sutter instruments Co. USA Model P.97). The odorants were delivered for duration of 1s at a rate of 1l/min switched by a solenoid valve controlled by pCLAMP and digitizer (1440A, Molecular Devices). Hexanol, Nonanol, Octanoic acid, Geraniol (Sigma Aldrich) were used for odor stimulation. The data was digitized at 10 KHz using a digitizer (1440A, Molecular Devices) and analyzed using a custom program written in MATLAB (Mathworks, Natick, MA).

## Behavior

Bees were collected from the hive in the evening. After cooling they were harnessed in plastic tubes with adhesive tape on their neck region. They were fed 1M sucrose solution (Sigma Aldrich) ad libitum and stored in a humid, dark chamber. Next morning the ones that showed robust proboscis extension response (PER) were selected for training. (Bitterman et al. 1983; Giurfa and Sandoz 2012; Matsumoto et al. 2012; Menzel et al. 2001)

The bees were placed in front of an odor delivery system with an air suction vent placed behind. They were left for 25 secs before stimulus application to reduce contextual stimuli. The odor delivery was controlled by a computer. The bees were presented with 4 sec odor stimulation with 2-Octanol (Sigma Aldrich). After the 3rd second the bees were given sucrose stimulation for 3 secs. First, the antennae of the bees were touched with a toothpick dipped in 1M sucrose solution and later they were allowed to feed on it by touching the proboscis. The onset of odor and the time for sucrose stimulation were indicated by green and red lights placed such that the bees cannot see them. After training, the bees were again left for 25 secs before being removed. Equal numbers of bees (20 in each group) were trained with 30 sec, 3 min and 10 min inter-trial intervals (ITI). To test odor discrimination two groups of bees (20 in each) were trained with Hexanol and Geraniol (Sigma Aldrich) as conditioned odor (CS) and novel odor (Nod) and vice versa. The above odors were chosen as they were found to be discriminated well from each other in *Apis mellifera* (Smith and Menzel 1989). The bees were tested 60 mins after the acquisition. Bees that did not have robust PER were not included in the testing. Cochran’s Q was used to check the effect of trials on the acquisition. Data was analyzed using program in Python.

## Results

### Tracts of the antennal nerve innervating the antennal lobe

In *Apis dorsata* too ORNs innervate the antennal lobe through the four tracts (T1-4) of the antennal nerve (Fig 1). Tracts T5 and T6 bypass the AL and enter the dorsal lobe and the protocerebrum respectively. The T1 tract innervated around 70 dorsal anterior glomeruli in the three counted samples, T2 tract was found to have two tracts T2-1 that innervated 1 glomerulus and T2-2 that innervated 6 medial glomeruli. T3 tract further divided into T3a, b and c tracts. It innervated around 70-80 ventro-posterior glomeruli. T3b glomeruli are located separately from the rest, caudal-dorsal to the T1 tract, they are smaller in size compared to the other glomeruli and the ORNs innervate along with the periphery, the entire core region and hence were termed ‘lobule’ by Arnold et al 1985 (Krischner et al. 2006). We counted around 12-16 T3b glomeruli. T4 tract innervated 7 posterior glomeruli. The total glomerular counts were 165, 164 and 165 in the three individuals that were counted. Table 1 provides the glomerular count of *Apis dorsata* and Table 2 has the count for *Apis mellifera* from different studies

**Fig. 1a-f.**
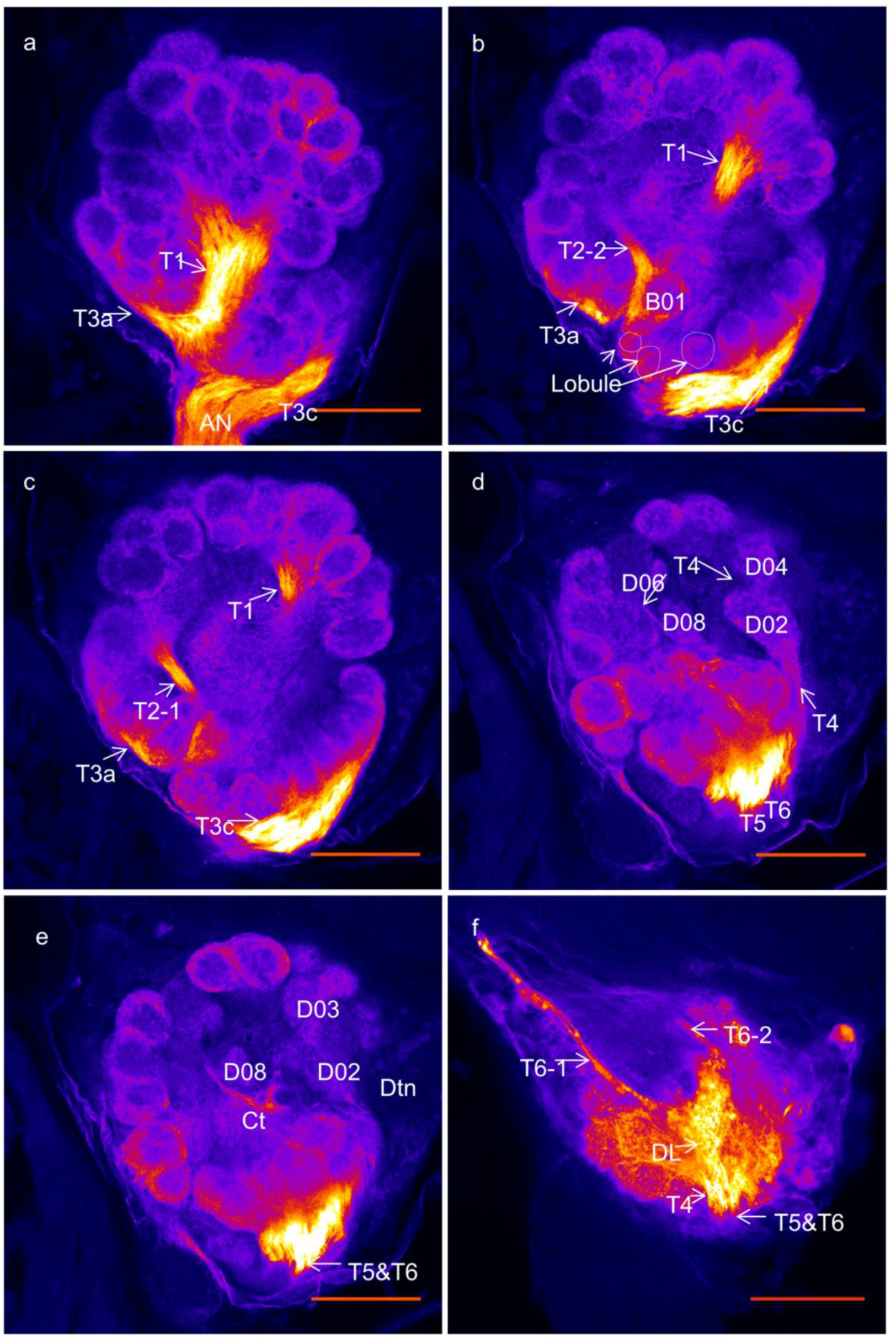
Optical sections of AL showing the tracts T1-6 a. T1 tract that innervates dorsal anterior glomeruli and T3a and T3c that innervate ventral posterior glomeruli are marked with arrows. b. T2-2 can be seen innervating one glomerulus. T3b glomeruli termed ‘lobule’ due to their small size and innervation pattern are circled. c. T2-1 that innervates 6 medial glomeruli can be seen. d. T4 tract that further divides in to 7 distinct tracts innervating 7 most posterior glomeruli is visible. e. The crescent tract (ct) that originates between D03 and D02 and acts as a land mark in glomerular identification can be seen clearly. D08 lies rostral to the ct. f. T5 is seen ending in the dorsal lobe and T6 is seen projecting towards the protocerebrum. Dtn, deuteoneurons. Scale bar = 100μm.

**Table.1.** Total glomerular count and count of glomeruli innervated by T1-T4 tracts in *Apis dorsata*. a,b,c represent the three counted samples whose total number and number of glomeruli innervated by tracks T1-4 is given.

**Table.2.** Glomerular count of *Apis mellifera* from existing studies. Total count and count of glomeruli innervated by different tracts of the AN in *Apis mellifera* according to different studies.

### The arrangement of the glomeruli of *Apis dorsata* antennal lobe

All the primary glomeruli were identified and were found to be homologous to those of *Apis mellifera* (Arnold et al. 1985; Flanagan and Mercer, 1989; Galizia et al. 1999). In the case of the secondary glomeruli, slight variations were found only in the position. Similar glomerular characteristics as described by Arnold et al, Flanagan and Mercer in *Apis mellifera* were also observed in *Apis dorsata*, (Fig. 2). A42, A33 and A17 are three big primary glomeruli in the anterior part of the AL which were easily identified. A4, A7, A11 and A22 are four primary glomeruli present at the lateral passage. In most of the cases, though A44 is the biggest T1 glomerulus we found A39, A40, A46, and A70 also to be big in size compared to the others. A14, A41 and A58 were easily identifiable rostral T1 glomeruli due to their closeness to T4 cluster. Among the T2 glomeruli, B01 was easily identified due to its inward orientation and the remaining ones could be identified due to their proximity to B01. In the T3 cluster, C23 is located at the origin of the T1 tract and was identified unambiguously. T3b glomeruli varied in number in the three counted samples as observed in *Apis mellifera*. T4 cluster glomeruli were large in size with ORN innervation in the center too and located posteriorly. The glomerulus D08 reported by Krischner et al. could be clearly identified in all three of the counted ALs (Fig. 2e). The Digital atlas of the glomeruli of *Apis dorsata* based on Galizia et al 1999 is available as electronic supplementary material, ESM_1.

**Fig. 2a-f.**
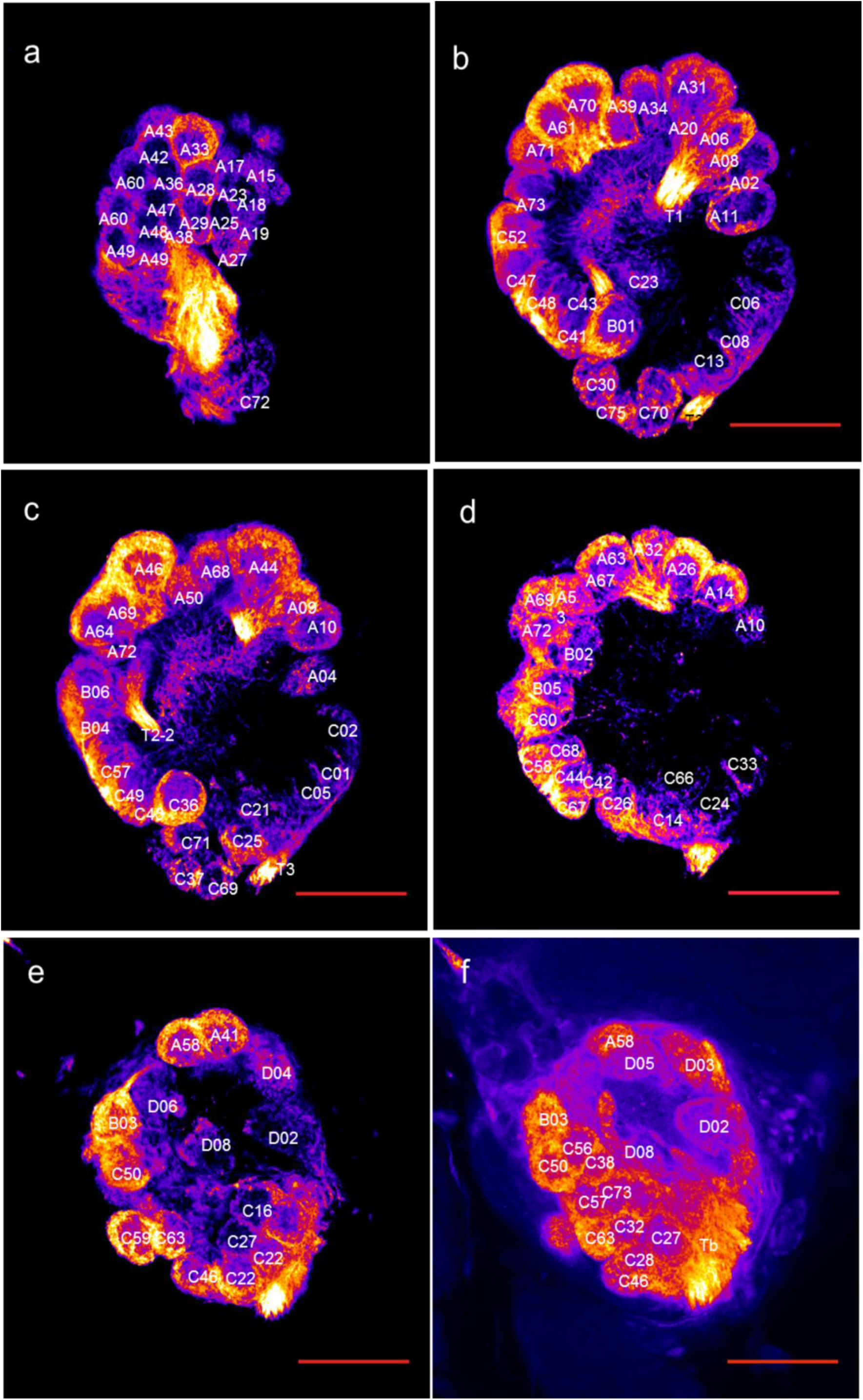
Optical sections of the AL at different depths showing various primary and secondary glomeruli a. The three primary glomeruli A33, A32, A17 can be seen. Glomeruli A49 and A60 are present in duplicate. b. Primary, invariant glomerulus C23 is present at origin of T1 and T2 tracts. c. A44 the largest T1 glomerulus and T3b glomeruli C71, C37, C69 can be seen. It can be noted that T3b glomeruli differ from the rest in their size and innervation pattern. d. Primary glomeruli B05 and B02, the biggest T2 glomerulus are visible. e. A41 a big, primary glomerulus closest to the T4 cluster is seen. f. T4 glomeruli D04, D02, D08 and D06 can be seen. Scale bar= 100μm.

Some glomeruli (around 10 in an individual) were missing in some and few glomeruli (around 5) were found in double in some. For example in sample A glomeruli A5, A35, A37, A40, A51, A55, A66, and C23 were missing whereas A49, A60, A62, C22, C66 and C34 were found in double. In sample A16, A54, A59, A71 and C39 were missing and A27, A62, C6, C45, C52 and C55 were double. In *Apis mellifera* too similar phenomenon has been reported by Arnold et al 1985 and Galizia et al 1999. Individuals collected from different hives showed more variation in glomerular position and size than individuals from the same hive. In Fig. 3, a and b are from the two sides (left and right) of the same individual and, c and d are of individuals from different hives. It can be noted that there are slight variations in the position of glomeruli. When both the ALs of an individual were filled and studied, the glomeruli in right and left AL were found to be identical in number and spatial arrangement suggesting that the variability observed between individuals are real and not an artifact of tissue processing or imaging (Fig. 3a and b).

**Fig. 3a-d.**
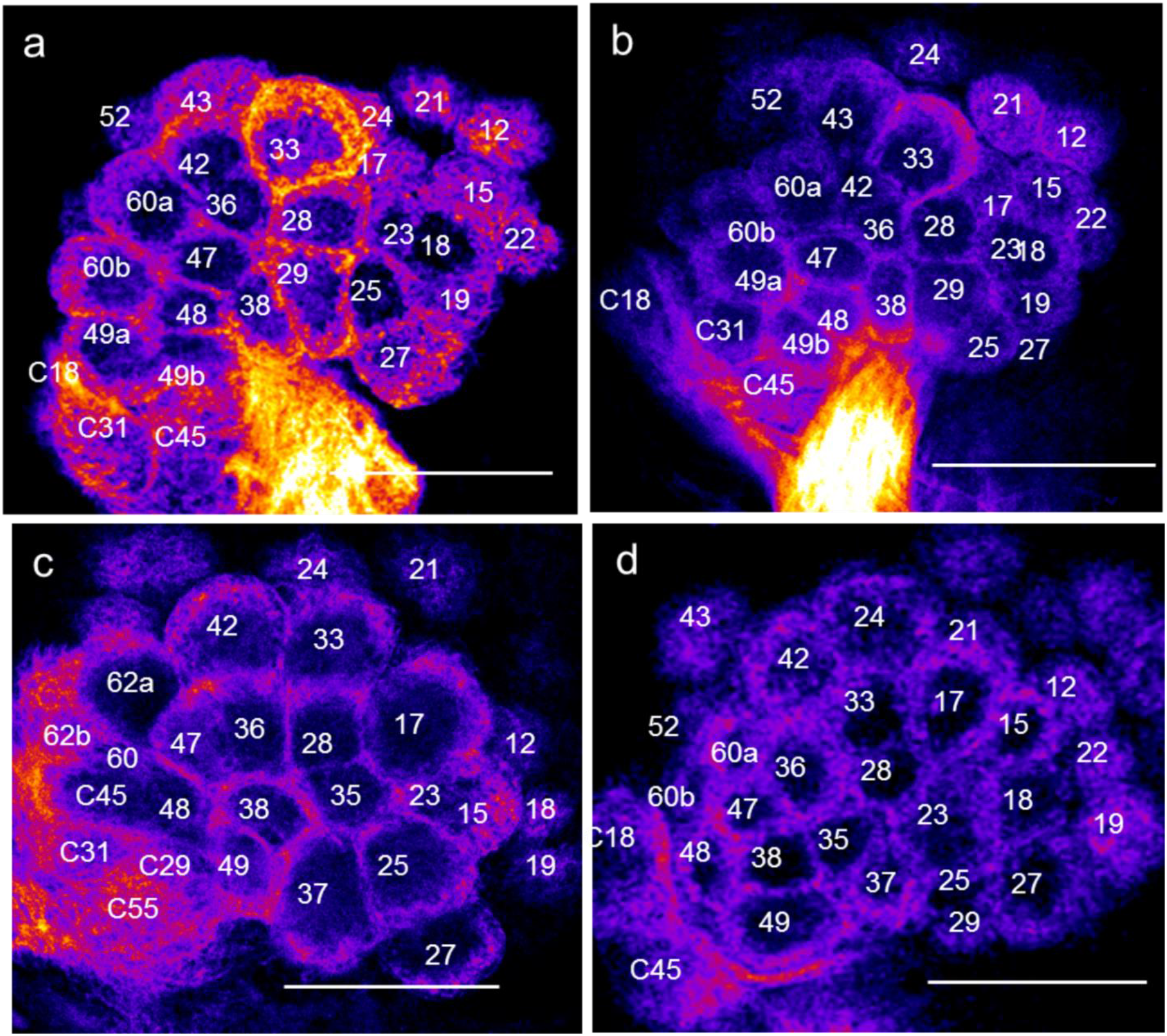
Inter-individual differences in glomerular organization a and b. optical sections of the right and the left ALs of an individual. It can be noted that they are identical in glomerular arrangement. Glomeruli A35 and A37 are missing and A60 and A49 are double in both a and b. c and d. ALs of individuals from different hives. Slight variations in position of the glomeruli between a and b, c and d can be seen. In c it can be noted that glomerulus A62 is double and in d A60 is double. Scale bar= 100μm.

A 3D model of the AL of *Apis dorsata* is shown in Fig. 4. Each division was labeled using a different color (Krischner et al. 2006). T1 is colored red, T2 is yellow, T3a light blue, T3b purple, T3c dark blue and T4 is green. A 3D animation of the antennal lobe is available as Electronic supplementary material, ESM_2.

**Fig. 4a-d.**
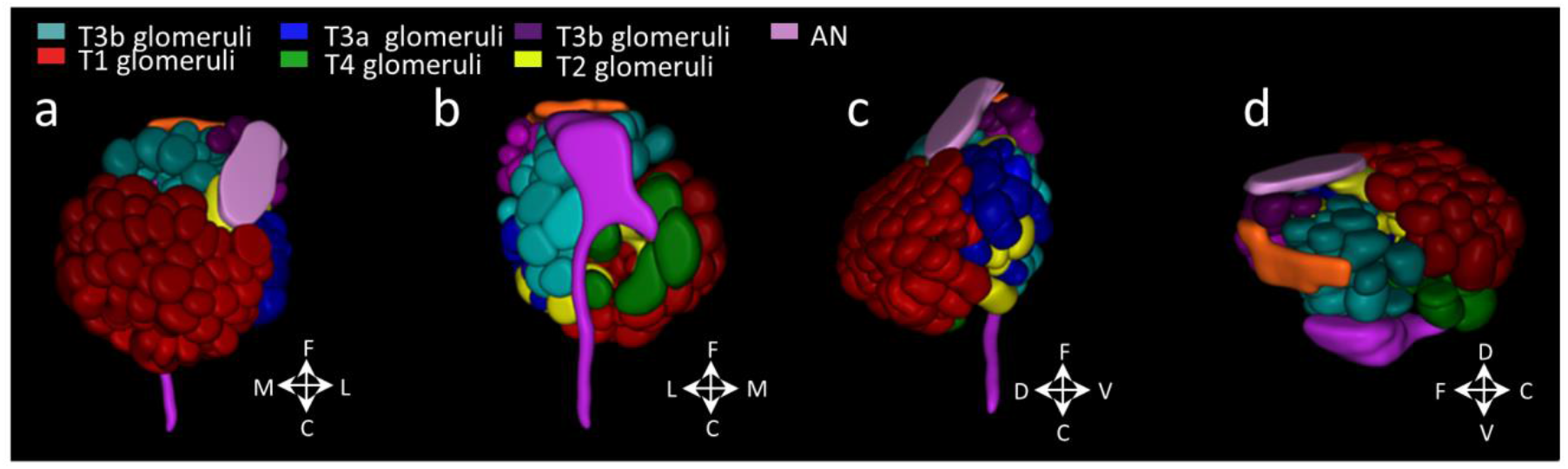
3D model of the antennal lobe showing the arrangement of the glomerular clusters a. Dorsal view showing the antennal nerve entering the antennal lobe. b. Ventral view showing T6 nerve. c. Side view showing the lateral side of the antennal lobe. d. Side view showing medial side of the antennal lobe.

### Circuit of the antennal lobe

Dextran fill from the antennal nerve shows ORN terminals in the periphery of the glomeruli. Immunohistochemistry revealed periphery of the glomeruli innervated by the ORNs is GABA negative whereas the inner core was GABA positive (Figs. 5a, b, and c). analogous to that reported in *Apis mellifera* (Arnold et al. 1985; Galizzia et al. 1999) Dextran injection in the AL makes the cell bodies of the antennal lobe visible (Fig. 5d) They are mostly located on the lateral side of the antennal lobe. Immunohistochemistry (Fig. 5e) shows that GABA positive cell clusters of putative LNs are in the lateral side of the AL (Fig. 5e).

**Fig. 5a-e.**
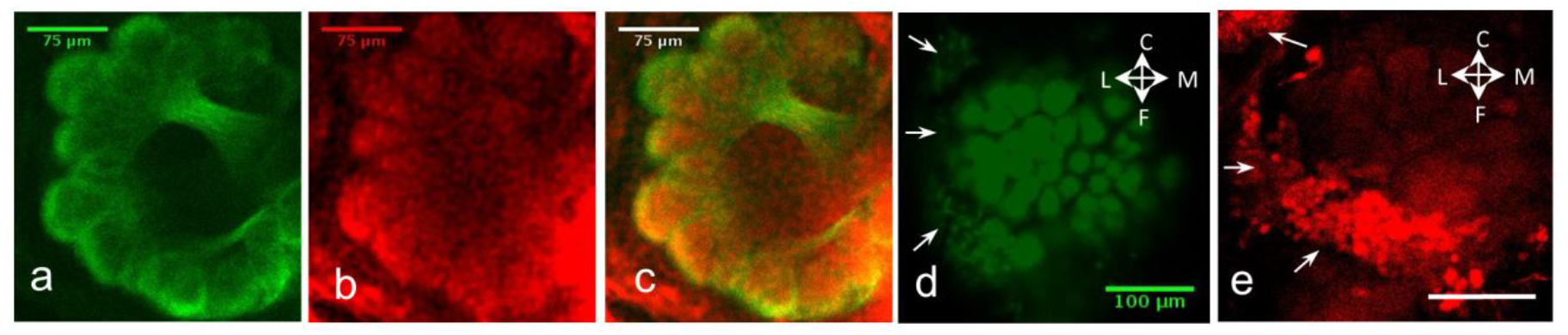
GABA positive regions innervate the inside of the glomeruli while the ORN fibers innervate the periphery a. The ORN innervations visualized by dextran fill from antennal nerve. b. Regions of GABA positivity inside the glomeruli. c. Merge of a and b. d. Cell bodies of the neurons of the AL are predominantly on the laterals side as revealed by dextran tracing from the AL. e. GABA positive cell clusters of the AL are on the lateral side. Scale bar = 100microns.

### Antenno-cerebral tracts of *Apis dorsata*

The anterograde mass fills of AL revealed that the efferent tracts of the AL were also similar to *Apis mellifera* (Mobbs 1982; Abel et al. 2001; Krischner et al. 2006; Muller et al. 2002; Rossler and Brill 2013; Zwaka et al. 2016). In *Apis dorsata* too axons of the PNs project to the protocerebrum via 5 Antenno-cerebral tracts, the medial (m-ACT), lateral (l-ACT) and mediolateral (ml-ACT) tracts (Fig. 6). The m-ACT travels dorsally to the mushroom body calyces where it sends out collaterals to the lip of both the calyces, and then it travels ventrally to the lateral horn (LH). The l-ACT leaves the AL and innervates the LH first and then projects to the Mushroom body calyces. The three ml-ACTs branch off from the m-ACT at different depths. ml-ACT-1 branches first from the m-ACT and innervates the LH. It does not have branches. mlACT-2 bifurcates into two, one branch goes to the LH and the other branch projects to the ring neuropil of the alpha lobe. ml-ACT-3 was found to be made up of a network of four tracts (Fig. 6) each subtract branches off from m-ACT at different positions and projects towards the lateral protocerebral lobe. ml-ACT-3a and ml-ACT-3b project towards the base of the calyx, ml-ACT-3c innervates the LH and ml-ACT-3d bends before branching into two branches which terminate on the l-ACT.

**Fig. 6a-b.**
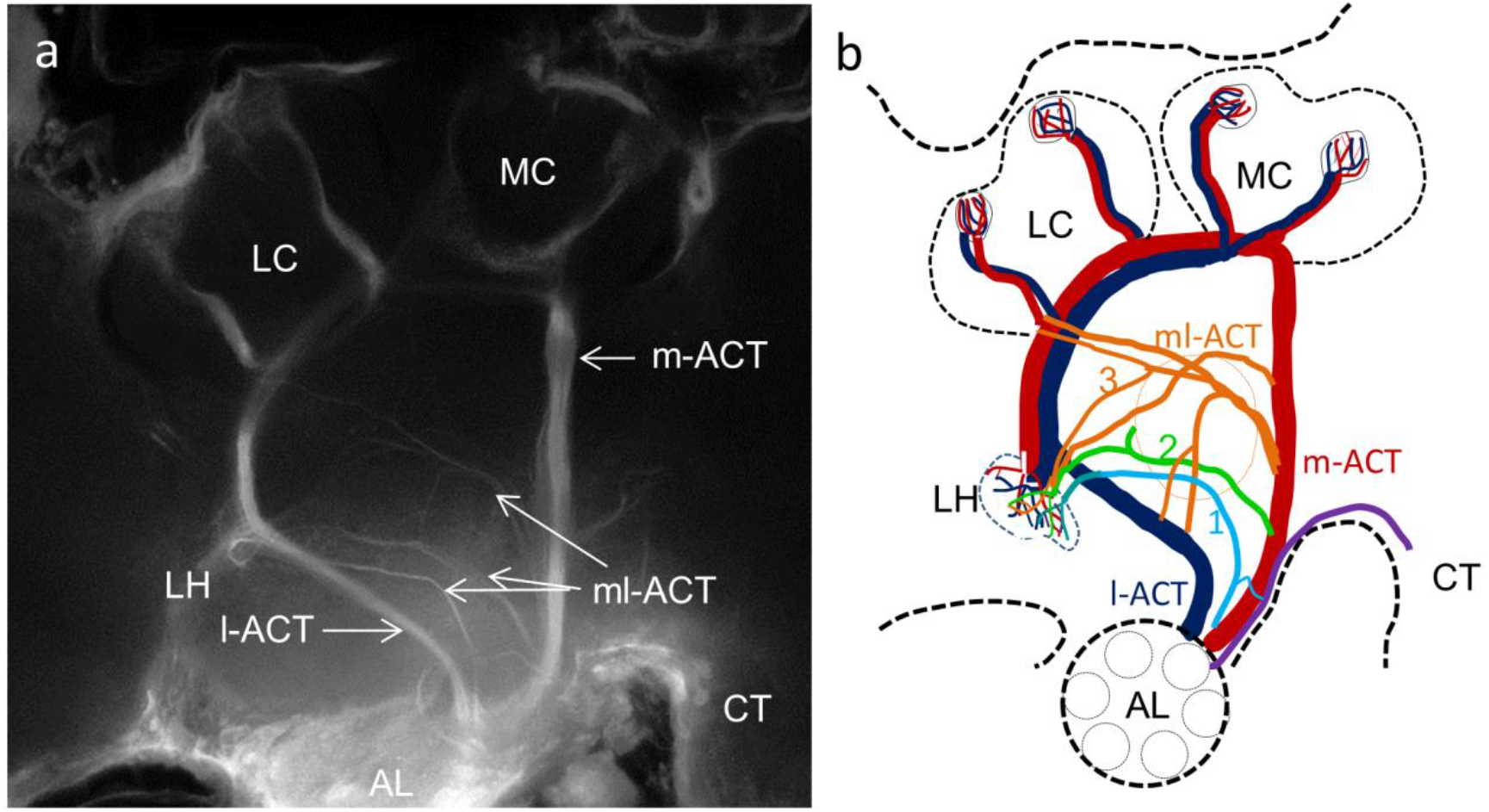
One odor evokes different types of temporally patterned responses in different cells and different odors evoke different types of temporally patterned responses in the same cell a. Responses of different neurons of the antennal lobe to a single odor (Hexanol 100%). b. Responses of a single cell to four different odors. The odor Hexanol was repeated at the end to show the invariability of odor responses.

A thin tract was found to innervate the contralateral AL similar to what has been reported in *Apis mellifera* by Arnold et al 1985.

### Neurons of the antennal lobe respond in odor-cell specific way

We recorded intracellularly from the AL interneurons to know their response properties. Cells had a baseline firing rate of 1 to 2 spikes per second and exhibited varied responses when presented with different odors. Some cells were inhibited by an odor. For example in Fig. 7a the cell was inhibited by Nonanol whereas Hexanol produced excitation. Different cells exhibited different temporally patterned responses to a single odor. Fig. 7b shows responses of different cells to the odor Hexanol. These cells often had a spontaneous spiking activity even when odor stimulus was not applied.

**Fig. 7a-b.**
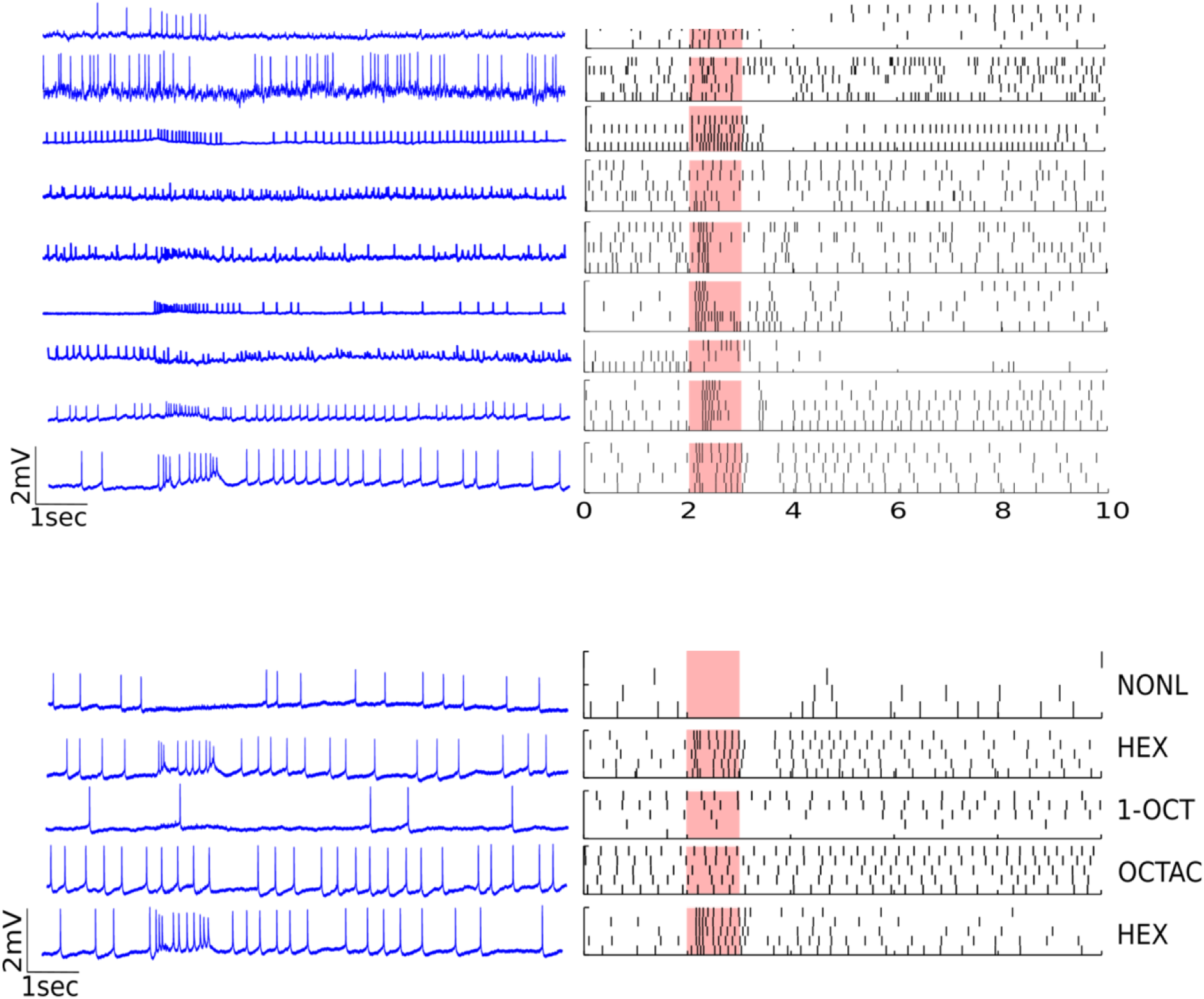
Projections of the AL efferent tracts a. An anterograde fill of AL showing the projections of all 5 PN tracts. The m-ACT projects first to the MB calyces then to the LH, the l-ACT innervates the LH first and later projects to the MB calyces. ml-ACTs branch off from the m-ACT at different depths and innervate the lateral protocerebral lobe. ml-ACT1 runs to the LH without branching. ml-ACT2 branches into two, one branch goes to the LH and other innervates around the alpha lobe. ml-ACT3 is made of 4 sub tracts. ml-ACT3-a and b project to the base of calyx, ml-ACT3-c innervates the LH and ml-ACT3-d is seen bending and terminating on the lACT. Some axons can be seen going to the contralateral Antennal Lobe (CT). b. Schematic showing the innervation pattern of the 5 tracts. LC lateral calyx; MC medial calyx; CT contralateral AL.

### Structure of the mushroom body

Dextran tracing from the calyx in a sequence of locations showed that axons that were filled and unfilled stay partitioned in the peduncle and the lobes (Fig. 8a, b). Kenyon cells with cell bodies in different regions in the calyx send their axons in parallel tracts in the peduncle and the alpha lobe. Thus they have a columnar arrangement and preserve neighborhoods and compartmentalization in the peduncle and lobes.

**Fig. 8a-d.**
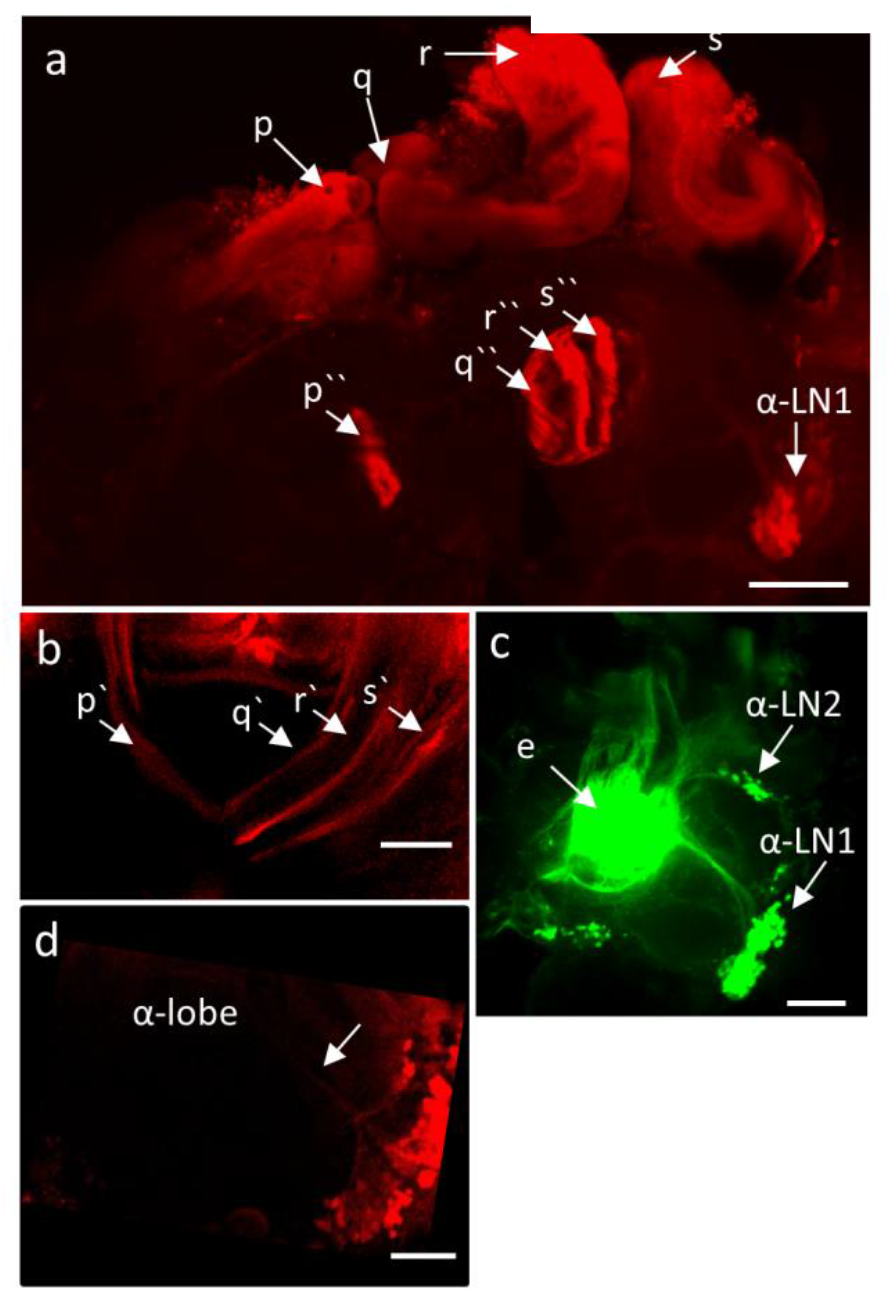
Mushroom body circuitry of *Apis dorsata* a and b. Dextran tracing shows that the Kenyon cells form a columnar arrangement that partitions the mushroom body into parallel compartments. p, q, r and s in a show locations of dextran injection in the calyx. p`, q`, r` and s` in b are the axons of the corresponding Kenyon cells visible in the peduncle and beta-lobe. p``, q``, r`` and s`` are the axons of the corresponding Kenyon cells visible in the α-lobe. a and c. the mushroom body extrinsic neurons α-LN1. In a they are filled from calyx and in c they are filled from alpha lobe. α-LN2 are another set of cells filled from alpha lobe. d. GABA positive cells can be seen sending neurites to the alpha lobe. Scale bar = 100microns.

### Extrinsic neurons of the mushroom body are restricted to the ipsilateral side

Dextran tracers injected in to the alpha lobe filled extrinsic neurons of the mushroom body which innervate calyx and cell bodies in the lateral protocerebrum (Fig. 8c). Dextran tracing from calyx also showed similar innervations to alpha lobe and cell body locations (Fig. 8a). In each case, around 35 cell bodies could be counted. No contralateral connections were observed when the dextran tracing was done from the calyx or the alpha lobe. Immunohistochemistry revealed GABA positive cells in similar locations with neurites leading to alpha lobe (Fig. 8d).

### *Apis dorsata* shows robust olfactory PER conditioning

We tested the amenability of *Apis dorsata* to olfactory conditioning. These bees had very low PER (less than 4%) to naive odor. After training using olfactory conditioning protocol (n=60) (Bitterman et al. 1983; Menzel 1990; Matsumoto et al. 2012) we found that odor evoked PER scores significantly increased up to 92% with trial number. They were amenable to both massed and spaced training and showed a significant increase in PER scores for 30 sec, 3 min and 10 min inter trail intervals (Fig. 9a) (Cochran’s Q values 46.9, 41.09, 49.2; df=6; p<0.001). They exhibited both short term and long term memory. They had retention of 57, 47 and 52 percent respectively one hour after conditioning for the 3 ITI’s and retention of 28, 33 and 42 percent 24 hours after conditioning (Fig. 9 b). They were able to distinguish a trained odor from a novel untrained odor. Bees trained with Octanol or Geraniol as the CS with 6 conditioning trials showed 0% response to novel odor (the untrained among the two) and 50% to CS one hour after conditioning (Fig. 9c).

**Fig. 9a-c.**
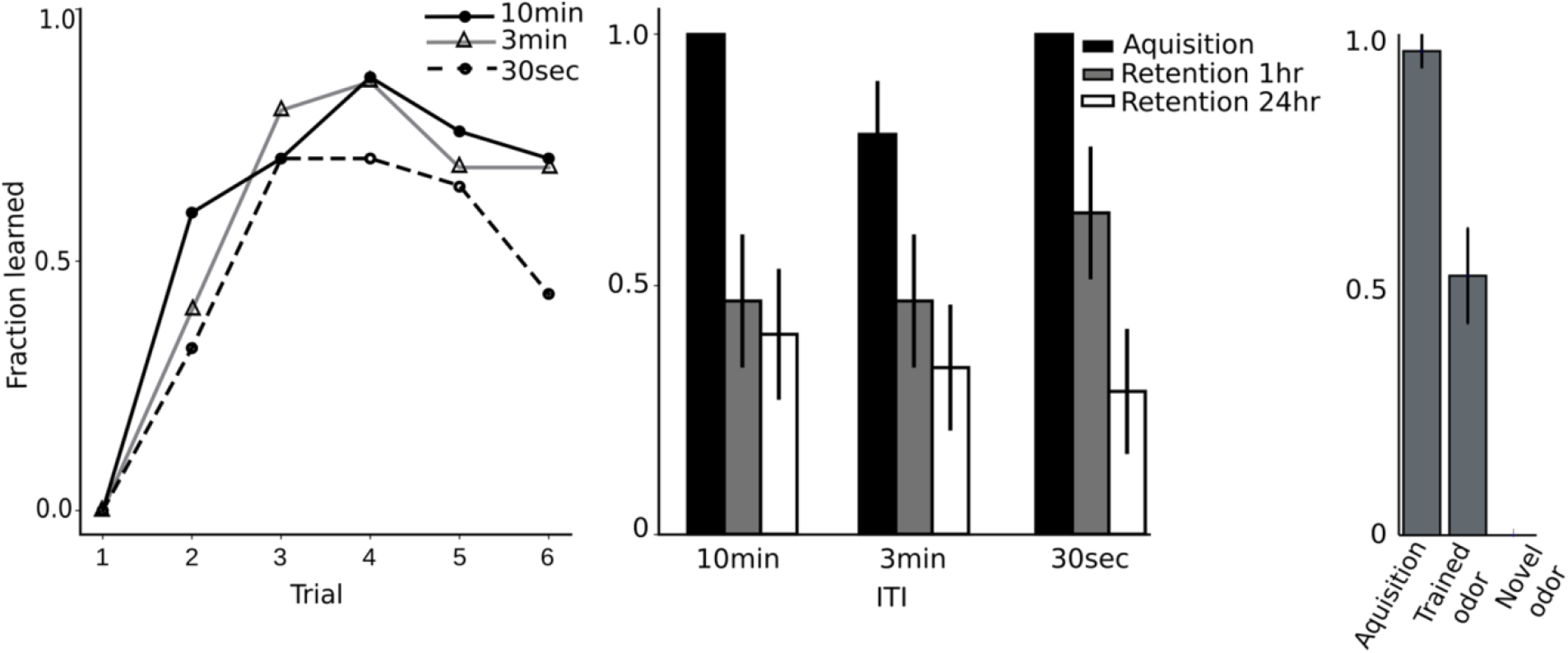
*Apis dorsata* learns and retains memory in PER olfactory conditioning paradigm a. The Learning curves for three different inter- trial intervals ranging from spaced (10mins) to massed (30sec). b. Memory tested at 1 hour and 24 hours compared to the acquisition at 6^th^ trial. c. Acquisition at 6^th^ trail and response after one hour to trained odor and untrained novel odor

## Discussion

*Apis dorsata dorsata* is an open nesting bee belonging to the genus Apis, subgenus Megapis and is widely distributed in India and southern Asia. In this study, we characterize the olfactory system of *Apis dorsata*. The sensilla on the antenna in this species have already been characterized (Neelima R.Kumar et al. 2014). The ORNs present in the antennal sensilla innervate the glomeruli of the ipsilateral AL through the AN. The arrangement of these tracts in space and thickness was very similar to *Apis mellifera* (Electronic supplementary material Fig. S 1)(Mobbs 1982) Based on the four AN tracts innervating them, in *Apis dorsata*, the glomeruli were divided into T1, T2, T3 and T4 glomeruli analogous to other *Apis* species. T1 glomeruli were numbered with a prefix A, T2 with B, T3 with C and T4 with D. When the glomeruli were named following the nomenclature applied by Galizia et al. (1999) there is one to one correspondence in the glomerular set indicating the close similarity of the two *Apis* species.

Krischner et al. (2006) had reported a novel glomerulus (compared to Galizzia et al. 1999) like structure innervated by T4 tract and named it D08 and reallocated D07 to T3 division and named it C73 as it was found to be innervated by T3 tract. We found a similar arrangement in *Apis dorsata*. Similar to the finding of Arnold et al 1985 and Galizzia et al 1999, we too found the ORN innervation only in the cortex of the glomeruli in all cases except in T4 and T3b glomeruli. We also found the ORN innervation to be non-GABAergic. However, the inner core was found to be GABAergic. We further need to establish which subsets of PNs and LNs are GABAergic. In the case of *Apis mellifera* 750 out of 4000 of the LNs were found to be GABAergic (Bicker 1986).

The glomerular arrangement and number were found to be similar to *Apis mellifera* in all the three counted samples. The glomerular count is a good constraint for searching for olfactory receptor genes as most of the glomeruli receive inputs from all receptor neuron expressing a single receptor type (Robertson and Wanner 2006). As expected from this line of argument, both *Apis florea* and Apis *mellifera* were found to have similar number of olfactory receptor (OR) genes (Karpe et al. 2016) that equals their glomerular counts. *Apis dorsata* might also have a similar number of OR genes as *Apis mellifera* and *Apis florea*. Though the number and arrangement of glomeruli is similar, the odor representation in the glomeruli cannot be assumed to be similar because the receptors types of the two species can be different. This is something to be tested. We have created the digital atlas and made it available online as Supplementary material. This can be used to make a functional atlas of odor representation. Digital atlas can also aid in the identification of glomeruli innervated by AL neurons while carrying out intracellular fills. Our AL filling showed that the PN axons leave the AL via the 5 ACTs. In *Apis mellifera* the m-ACT and l-ACT contain primarily axons of uniglomerular PNs (Bicker et al. 1993; Abel et al. 2001; Brandt et al. 2005)) whereas ml-ACT contain mostly axons of multiglomerular PNs (Fonta et al. 1993). Further l-ACT PNs belong to the T1 cluster and m-ACT PNs majority belong to the T3 cluster. These details are yet to be established in *Apis dorsata*. Therefore the computation that is posited to be carried out by this dual path arrangement of antenno-cerebral tracts in Apis *melliera* is unconfirmed as yet in *Apis dorsata*.

Bilateral connections in Apis.

In *Apis mellifera* it has been shown that the memory from one side can be transferred to the other side after three hours. However in the dextran tracings from antenna, AL, calyx of the mushroom body and the alpha lobe we could detect connections to the contralateral side only from the antennal lobe. Thus the mechanism of this reported transfer remains a mystery.

PER conditioning paradigm in *Apis mellifera* has been a versatile tool to investigate aspects of the classical conditioning. The difference in the nature of memory, when trained under massed and spaced protocols, has been used to understand the mechanisms underlying learning and memory (Tully 2005). Thus the ability to train *Apis dorsata* in these protocols effectively opens up the possibility of using this as a model system analogous to *Apis mellifera* to address such issues.

## Summary

A prominent difference between the four species of honey bees is their nesting habit. Among these the two that have been investigated the most are A*pis mellifera* and *Apis dorsata*. In them *Apis mellifera* is cavity nesting and *Apis dorsata* is open nesting. There are no prominent differences in the ORN tracts, glomerular counts and their position, the antenno-cerebral tracts or the arrangement further upstream in MB and their extrinsic neurons that enable us to distinguish the two. Moreover the *Apis dorsata* also behaves similarly to *Apis mellifera* in the PER conditioning assay.

## Acknowledgement

We would like to thank Uttam Krishna Sharma for his support in procuring honeybees and Shilpi Singh for her support in carrying out electrophysiology. We also thank Prasad Miriyala (Central Instruments Laboratory) and Nalini Manthapuram (Centre for Nano Technology) for their support in confocal imaging. We are grateful to UPE scheme of University Grants Commission, India and DST Purse for providing funding to University of Hyderabad.

**Figure S 1** Sagital view of the Antennal lobe showing the tracts T1 and T4 tracts can be seen innervating the glomeruli whereas T6 tract can be seen bypassing the AL and entering the protocerebrum.

## ESM_1

A digital atlas of the glomeruli of *Apis dorsata* based on the atlas of *Apis mellifera* made available by Galizzia et al 1999

## ESM_2

An animation of the 3D reconstruction of the antennal lobe of Apis dorsata. Glomeruli innervated by different tracts and the antennal nerve are labelled in different colors.

